# Multi-tissue Multi-omics Nutrigenomics Indicates Context-specific Effects of DHA on Rat Brain

**DOI:** 10.1101/2020.09.02.280289

**Authors:** Guanglin Zhang, Qingying Meng, Montgomery Blencowe, Agrawal Rahul, Fernando Gomez-Pinilla, Xia Yang

**Author notes:** Correspondence: Xia Yang, Ph.D., Department of Integrative Biology and Physiology, University of California Los Angeles, Los Angeles, CA 90095.

## Abstract

**Scope:** We explored the influence of DHA on cardiometabolic and cognitive phenotypes, and multiomic alterations in the brain under two metabolic conditions to understand context-specific nutritional effects.

**Methods and Results:** Rats were randomly assigned to a DHA-rich or a control chow diet while drinking water or high fructose solution, followed by profiling of metabolic and cognitive phenotypes and the transcriptome and DNA methylome of the hypothalamus and hippocampus. DHA reduced serum triglyceride and improved insulin resistance and memory exclusively in the fructose-consuming rats. In hippocampus, DHA affected genes related to synapse functions in the chow group but immune functions in the fructose group; in hypothalamus, DHA altered immune pathways in the chow group but metabolic pathways in the fructose group. Network modeling revealed context-specific regulators of DHA effects, including *Klf4* and *Dusp1* for chow condition and *Lum, Fn1*, and *Col1a1* for fructose condition in hippocampus, as well as *Cyr61, JunB, Ier2*, and *Pitx2* under chow condition and *Hcar1, Cdh1*, and *Osr1* under fructose condition in hypothalamus.

**Conclusion:** DHA exhibits differential influence on epigenetic loci, genes, pathways, and metabolic and cognitive phenotypes under different dietary contexts, supporting population stratification in DHA studies to achieve precision nutrition.

## 1. Introduction

Long-chain polyunsaturated fatty acids (PUFAs), particularly omega-3 (n-3) PUFAs, have been indicated to play important roles in various aspects of human health ^[1]^. Among the abundant n-3 PUFAs, docosahexaenoic acid (DHA; 22:6 n-3),in particular, has been extensively investigated. n-3 PUFAs may be beneficial for a broad range of disorders affecting both peripheral metabolism and brain functions, including obesity, hypertension, type 2 diabetes (T2D), insulin resistance, coronary heart disease (CHD), cardiovascular disease (CVD), cognitive disorders, and traumatic brain injury ^[2-5]^. Our previous research uncovered that rats fed with n-3 diet counteracted both metabolic and cognitive deficits induced by high fructose consumption ^[6]^.

Despite the numerous positive reports on the potential beneficial effects of n-3 PUFAs, controversies have also arisen. Multiple studies suggested that n-3 PUFAs have a favorable effect on plasma triglyceride levels but no effect on cholesterol, glycemic, insulin or insulin resistance in T2D or metabolic syndrome patients ^[7]^. Similarly, lack of effects has also been reported for the cardiovascular benefits of n-3 PUFAs ^[8-12]^.

We hypothesize that the conflicting findings may be a result of context-specific effects of DHA, that is, the benefits of DHA only manifest under specific pathophysiological conditions. Here we aim to employ high-throughput genomic and systems biology approaches to thoroughly investigate the phenotypic, transcriptomic, and epigenetic changes induced by DHA, a commonly studied n-3 PUFA, under different dietary contexts. Our results support profound context-specific alterations in genes, pathways, and epigenetic sites in individual brain regions affected by DHA despite certain similarities between contexts, thereby offering molecular insights into the differential activities of DHA that are dependent upon the physiological states of the host. These findings offer insights into the controversies observed in epidemiological and experimental studies regarding the benefits or lack of benefits of n-3 PUFA and support the need for more thorough future studies of PUFAs stratified by the metabolic conditions of the study subjects.

## 2. Experimental Section

### Animals

Two months old male Sprague-Dawley rats (Charles River Laboratories, Inc., MA, USA) were randomly assigned to omega-3 fatty acid diet rich in DHA (n = 8), or a control chow diet (#5001, LabDiet, St. Louis, MO) without omega-3 fatty acid supplementation (n = 8). The rats were singly housed with free access to drinking water or 15% fructose at room temperature (22-24°C) with 12h light/dark cycle. Metabolic phenotypes including serum levels of insulin, glucose, and triglycerides, and insulin resistance index (fasting glucose [mg/dl] × fasting insulin [ng/ml] / 16.31) were examined. Rats were trained in the Barnes maze device for 5 days before diet treatment, followed by memory retention test after 6 weeks of treatment. Then rats were sacrificed, and hypothalamus and hippocampus tissues were dissected out, flash frozen, and stored at -70°C.

### Dosage information

Rats were fed n-3 PUFA rich in DHA (1.2% of DHA, Nordic Naturals, Inc., CA, USA) at final dose of 620 mg/kg body weight which falls within common dose ranges ^[13]^ used in animals. According to the conversion guidance ^[14]^, the corresponding human equivalent dose is 100 mg/kg, which is within the dose range used in human clinical trials ^[15]^.

### RNA Sequencing (RNA-Seq) and Data Analysis

Total RNA was extracted from hypothalamus and hippocampus (n = 4 per dietary group per tissue; total 32 samples) using an All-Prep DNA/RNA Mini Kit (QIAGEN GmbH, Hilden, Germany). Sample size was based on previous RNA-Seq studies in which findings were validated using qPCR and gene perturbation experiments ^[6, 16, 17]^. Quantity and quality of RNA were checked using Qubit 2.0 Fluorometer (Life Technologies, NY, USA) and Bioanalyzer 2100 (Agilent Technologies, CA, USA). Two hypothalamus samples from the fructose group and DHA group failed standard quality control and were removed from the analysis. RNA-Seq libraries were prepared according to the standard Illumina protocol. Sequencing was performed in 100bp paired-end mode on HiSeq 2000 (Illumina Inc, CA, USA). RNA-Seq analysis was performed using the Tuxedo package as described previously ^[6]^. Genes and transcripts showing differential expression or alternative splicing at p < 0.01 in each brain region were defined as a gene “signature” for further integrative analyses. DEGs were assessed for enrichment of pathways using the KOBAS web server ^[18]^. Pathways at FDR < 5% were considered significant. RNA-Seq data was deposited to Gene Expression Omnibus (GEO) under accession numbers GSE59918 (both brain regions from control and fructose) and GSE64815 (both brain regions from fructose with DHA diets), and GSE89176 (both brain regions from the DHA diet).

### Network analysis of DHA DEGs

To investigate the gene-gene regulations among the DHA DEGs and identify potential regulators, we mapped the DEGs to previously constructed Bayesian network of brain tissues, as described in our previous study ^[6, 19]^ and used the weighted key driver analysis (wKDA) in Mergeomics ^[20]^ to identify key regulatory genes of the DHA DEGs from each tissue. Key Drivers (KDs) are defined as the network genes whose network neighboring genes were significantly enriched for DHA DEGs based on a Chi-square like statistic ^[20]^. Network genes reaching FDR < 5% were considered as potential KDs. The gene subnetworks of KDs were visualized using Cytoscape ^[21]^.

### Reduced Representation Bisulfite Sequencing (RRBS) of DNA Methylome

RRBS libraries of DNA samples (n = 4 per treatment group per brain region; total 32 samples) were prepared as described previously ^[6]^. One hypothalamus sample from the fructose+DHA group was removed due to low quality. Loci with methylation levels > 25% between groups and FDR < 5% were considered as differentially methylated loci (DMLs). DMLs adjacent genes (within 10kb) were assessed for enrichment of pathways using the KOBAS web server ^[18]^. Pathways at FDR < 5% were considered significant. RRBS data was deposited to GEO under accession numbers GSE59893 (both brain regions from control and fructose), GSE64816 (both brain regions from fructose with DHA diets), and GSE89176 (both regions from the DHA diet).

## 3. Results

### 3.1. Effect of DHA supplementation on metabolic and behavioral phenotypes

Under the chow diet condition, DHA supplementation did not significantly alter metabolic phenotypes including plasma triglycerides, glucose, insulin level and insulin resistance index (Figure 1A-D), or the memory phenotype measured by latency time in the Barnes maze test (Figure 1E), although there were non-significant trends toward decreased triglycerides and latency time. By contrast, when rats were fed a 15% high fructose diet to induce metabolic syndrome, DHA supplementation significantly improved insulin resistance index in addition to reducing triglycerides and latency time in memory retention ^[6]^ (Figure 1), indicating beneficial effects of DHA under a metabolically challenged condition.

**Figure 1.**
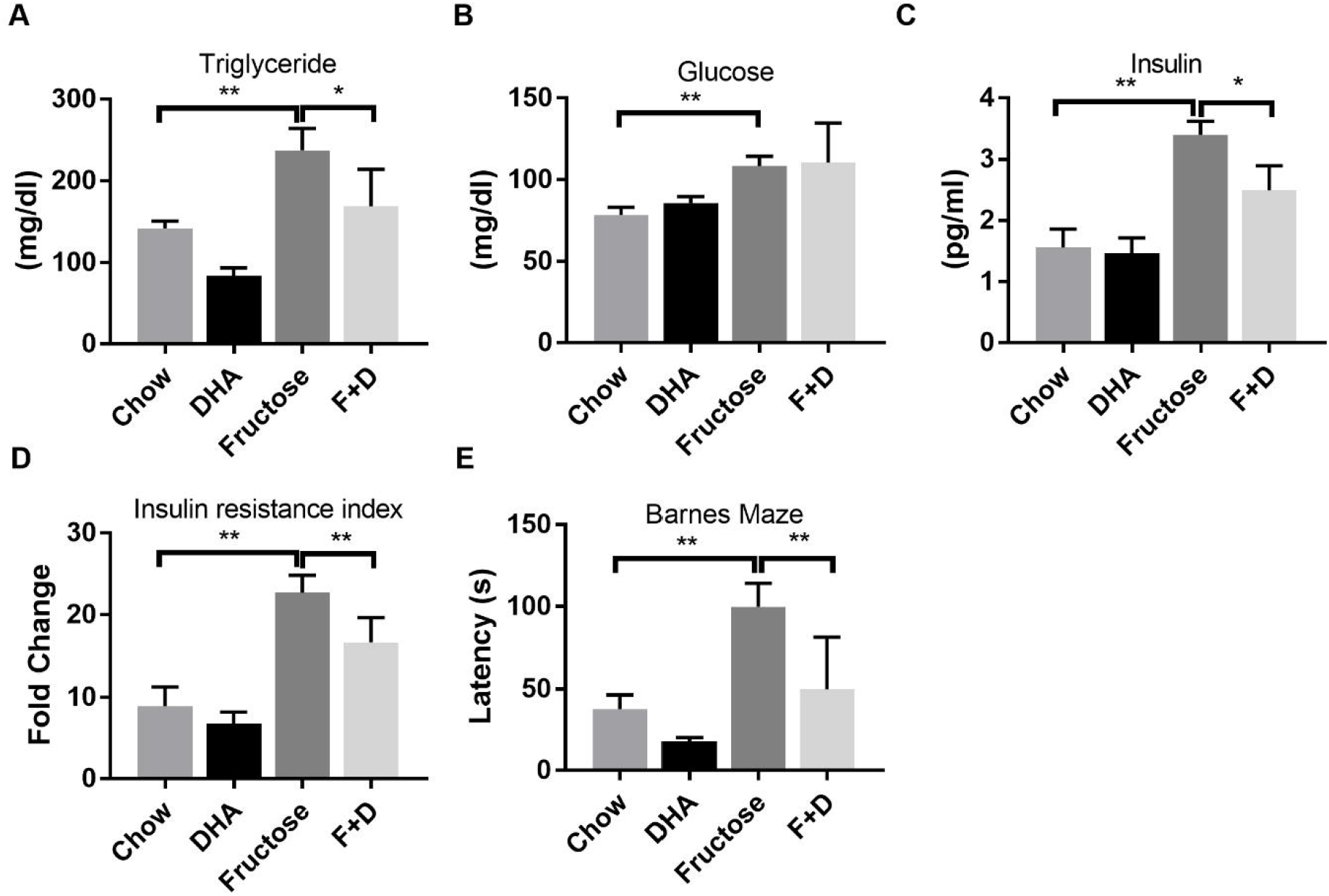
Changes in metabolic and behavior phenotypes in response to DHA supplementation under physiological and fructose-induced metabolic syndrome contexts. A) Serum triglycerides. B) Blood glucose. C) Insulin. D) Insulin resistance index. E) Latency time in the memory retention probe in the Barnes Maze test. * p < 0.05, ** p < 0.01 by ANOVA with Sidak test. Error bars in the plots are standard errors. N = 8/group. F + D: Fructose + DHA. Data of fructose and F+D from our previous study was used as comparison ^[6]^.

### 3.2. Transcriptomic alterations in rat hippocampus and hypothalamus induced by DHA supplementation to a chow diet

Despite a lack of significant phenotypic changes after DHA supplementation on a chow diet background, through RNA-Seq, we identified 141 and 388 DEGs, 86 and 252 differentially expressed transcripts, and 58 and 85 genes showing alternative exon usage in hippocampus and hypothalamus, respectively (Table S1). Overall, 214 unique hippocampal genes and 523 hypothalamic genes (including genes, transcripts, and alternative splicing) were defined as DHA DEGs, with 53 genes shared by the two tissues. The expression pattern changes between control and DHA groups are stronger in the hypothalamus (Figure 2B) than in the hippocampus (Figure 2A).

**Figure 2.**
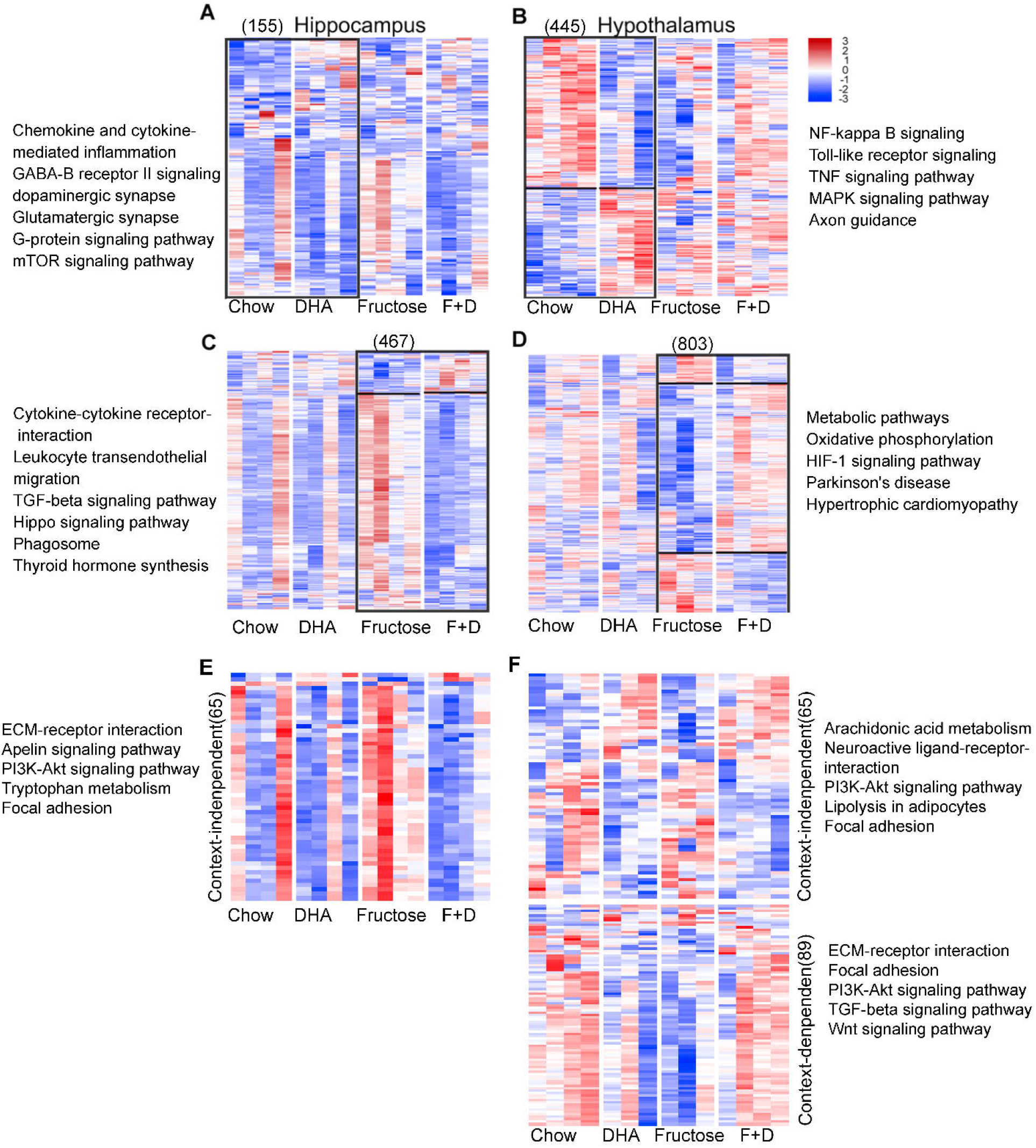
Heatmap of gene expression changes and enriched pathways under DHA supplementation with or without fructose treatment in hippocampus and hypothalamus. Unique DEGs affected by DHA in hippocampus (A) and hypothalamus (B) in chow diet background. Unique DEGs affected by DHA in hippocampus (C) and hypothalamus (D) when consuming fructose. Common DEGs affected by DHA in both chow and fructose backgrounds in hippocampus (E) and hypothalamus (F) were further divided into context-independent DEGs (upper panels) and context-independent DEGs (lower panel). DEGs were determined by Tuxedo at P < 0.01. Top significantly enriched pathways (FDR < 5%) among DEGs were shown. Blue to red colors indicate low to high expression values. The outer frame in each plot denotes the group comparison from which DEGs were chosen. Numbers in bracket represented the numbers of DEGs plotted. As alternative splicing has no expression values, the number was not included in the graph. F + D: Fructose + DHA. Data of fructose and F+D from our previous study was used as comparison ^[6]^.

To understand the biological functions of the DEGs affected by DHA supplementation, we evaluated the enrichment of the DEGs for biological pathways. We found 74 and 58 significantly enriched pathways at FDR < 5% from the hippocampal and hypothalamic DEGs, respectively, with 21 pathways shared by both (Table S2). The shared pathways between tissues include ECM-receptor interaction, Focal adhesion, and signaling pathways for PI3K-Akt, integrin, Rap1, and Wnt. The pathways unique to hippocampus include those highly relevant to neurotransmitter synapse functions, cardiomyopathy, and lipid metabolism. The hypothalamus-specific pathways include numerous innate immunity pathways, Axon guidance and Hippo signaling pathway (Table S2). These results support that DHA induces molecular alterations in diverse functional processes without significant phenotypic manifestation under a physiological condition.

### 3.3. Comparison of DHA DEGs on a chow diet background with those on a high fructose diet

We compared the DHA DEGs identified above from the chow diet background, representing a physiological condition, with those identified from rats consuming 15% fructose solution, a state of metabolic syndrome ^[6]^. As shown in Table 1, although there were significant overlaps in the DHA DEGs identified from the two conditions, many DEGs were context specific – either unique to each condition or show opposite patterns of expression changes between conditions. In general, more unique genes were affected by DHA when animals were fed a high fructose diet than when fed a control diet: 467 (Figure 2C) vs. 155 (Figure 2A) in hippocampus; 803 (Figure 2D) vs. 445 (Figure 2B) in hypothalamus. The expression patterns of DEGs affected by DHA supplementation differed between the chow and fructose backgrounds. For example, many DHA DEGs identified on the fructose background showed little discernable alterations by DHA when on a chow background (Figure 2C, 2D).

**Table 1.** Overlap of DEGs and methylation loci affected by DHA on a chow diet vs fructose diet background. Enrichment p value was calculated using 2-sided Fisher’s exact test.

| Omics | Tissue | DHA vs.<br>control | DHA+Fructose<br>vs Fructose | Overlap | Fold<br>enrichment | Enrichment<br>P value |
| --- | --- | --- | --- | --- | --- | --- |
| Transcriptome | hippocampus | 214 | 544 | 70 | 10.5 | $p < 2.8e-52$ |
| Transcriptome | hypothalamus | 523 | 920 | 165 | 6.0 | $p < 7.5e-85$ |
| DNA methylome | hippocampus | 1777 | 1957 | 343 | 862.3 | 0 |
| DNA methylome | hypothalamus | 1191 | 1665 | 195 | 657.5 | 0 |

DHA DEGs unique to the chow condition were enriched for pathways related to inflammation, neuronal functions, and metabolism of linoleic acid and ether lipid, and mTOR signaling in hippocampus (Figure 2A, Table S2) and immune pathways in hypothalamus (Figure 2B, Table S2). Under the fructose condition, unique DEGs affected by DHA were enriched for immune pathways, Hippo signaling, and thyroid hormone synthesis in hippocampus (Figure 2C, Table S2) and oxidative phosphorylation pathway in hypothalamus (Figure 2D, Table S2).

Among the common DEGs affected by DHA on both the chow and the fructose backgrounds, we observed two distinct patterns – context-independent (i.e., up or downregulated in both conditions) and context-dependent (i.e., up in one condition but down in the other). In both hippocampus and hypothalamus, the context-independent DHA DEGs are enriched for pathways including ECM-receptor interaction, PI3K-Akt signaling pathway, and Focal adhesion, indicating their essential role in the effects of DHA (Figure 2E). For the context-dependent DEGs affected by DHA, genes involved TGF-beta signaling pathway and Wnt signaling were enriched in hypothalamus (Figure 2F).

Together, these results suggest that DHA supplementation not only affects certain consistent pathways but exerts unique effects on gene expression programs depending on the dietary and physiological background.

### 3.4. Identification of tissue-specific key drivers (KDs) and subnetworks of DHA DEGs

To explore the potential regulatory genes that mediate the action of DHA on the downstream targets under chow diet or fructose diet, we used a data-driven network analysis to capture gene-gene regulatory relations. In hippocampus, we identified 37 and 122 KDs whose subnetworks were enriched for DEGs affected by DHA on the chow diet and the fructose diet, respectively, at an FDR < 5%; in hypothalamus 68 KDs and 132 KDs were identified under chow diet and fructose diet. These results suggest more extensive gene network changes in both brain regions induced by DHA under the fructose condition compared to the chow diet (Figure 3A,3B).

**Figure 3.**
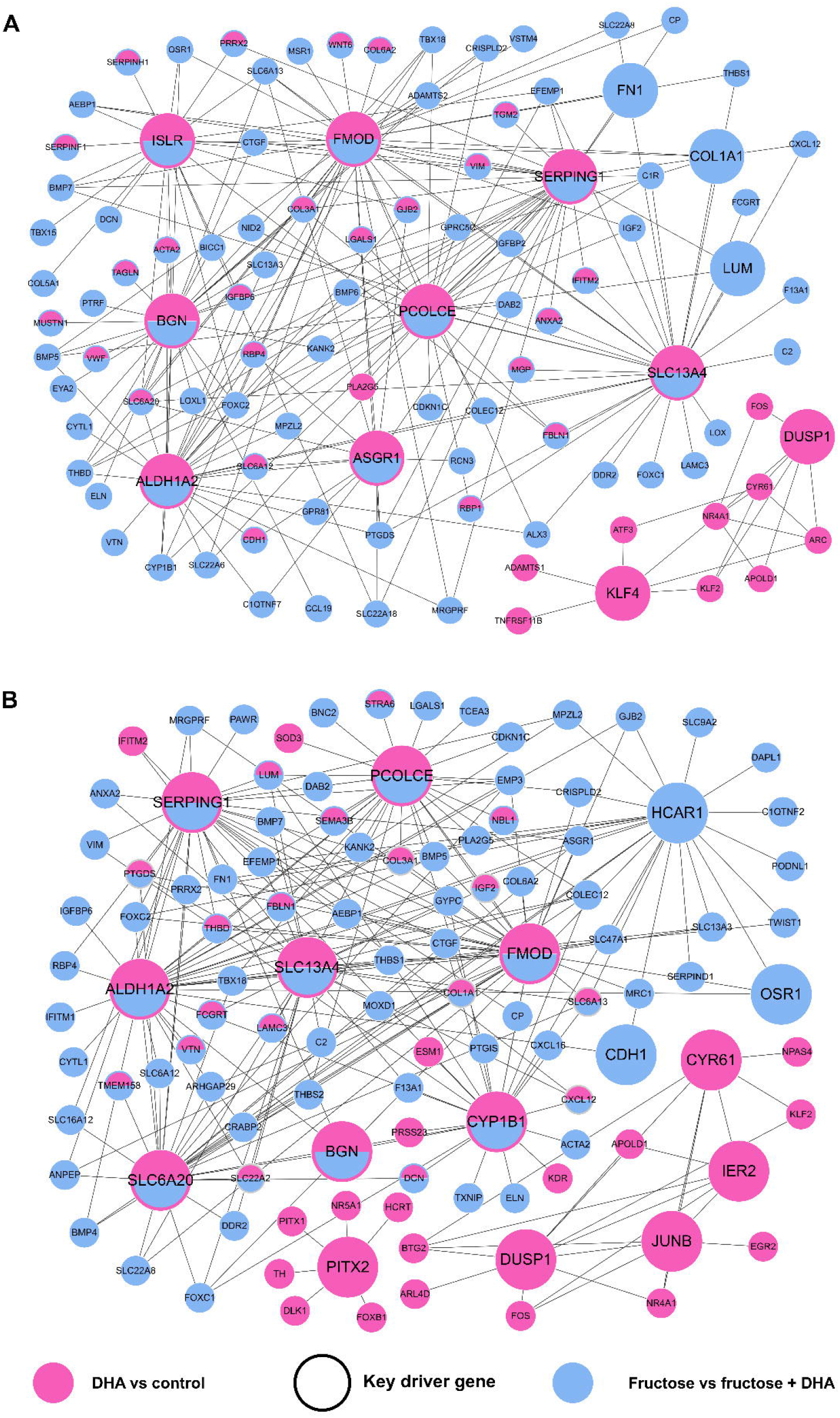
Gene subnetwork and top network key drivers of DEGs under different conditions affected by DHA in hippocampus (A) and hypothalamus (B). Larger nodes denote key driver; grey nodes denote genes not affected by DHA; green and yellow color represents hypothalamic and hippocampal DEGs, respectively.

Notably, the top KDs showed overlaps between chow and fructose conditions (Figure 3): *Fmod, Bgn, Serping1, Aldh1a2*, and *Pcolce*, mostly ECM genes, were top KDs for DEGs in both brain regions; *Slc13a4, Islr*, and *Asgr1* were consistent network KDs in hippocampus (Figure 3A); *Slc6a20* and *Cyp1b1* were shared KDs in hypothalamus (Figure 3B). Although these are shared KDs between chow and fructose conditions that are predicted to mediate the DHA effects on downstream genes, the DEGs surrounding these KDs in the gene networks are not necessarily the same between the chow and fructose conditions (pink versus blue nodes in Figure 3), suggesting differential gene regulation by these KDs depending on the interactions between DHA and the metabolic context.

In addition to these shared top KDs, we also found context-specific KDs of DEGs affected by DHA, including nine for chow condition (e.g., *Klf4* and *Dusp1*) and 95 for fructose condition (e.g., *Lum, Fn1, Col1a1*) in hippocampus, as well as 25 under chow condition (e.g., *Dusp1, Cyr61, JunB, Ier2, Pitx2*) and 27 for fructose condition (e.g., *Hcar1, Cdh1, Osr1*) in hypothalamus.

Collectively, ECM genes are the dominant and consistent type of KDs for DHA effects on the brain transcriptome, although other context-specific KDs such as *Klf4* and *Dusp1* for chow condition and *Hcar1* and *Osr1* for fructose condition also exist to regulate context-specific gene subnetworks. Many of these KDs are related to neuronal structural integrity and neural development and activity.

### 3.5. Context-specific DHA effects on the DNA Methylome

To explore the epigenetic mechanisms underlying the regulation of the transcriptome associated with DHA, we profiled the DNA methylome in both hippocampus and hypothalamus. We identified DMLs between DHA treated and untreated groups at FDR < 5%.

Comparison of the DHA-associated DMLs identified on the chow diet background with those from the fructose-fed conditions revealed both significant overlaps and DMLs unique to each condition (Table 1). Among the shared DMLs in both conditions, both context-independent (i.e., consistent changes by DHA regardless of chow or fructose background; upper panels) and context-dependent DMLs (i.e., opposite changes by DHA in chow vs. fructose conditions; lower panels) were identified (Figure 4E-F). Interestingly, when both DHA and fructose were consumed, the DNA methylation patterns of the context-dependent DMLs were normalized back to the patterns in the chow diet condition. Therefore, agreeing with the transcriptome level data, our DNA methylome data also support context-specific effects of DHA supplementation.

**Figure 4.**
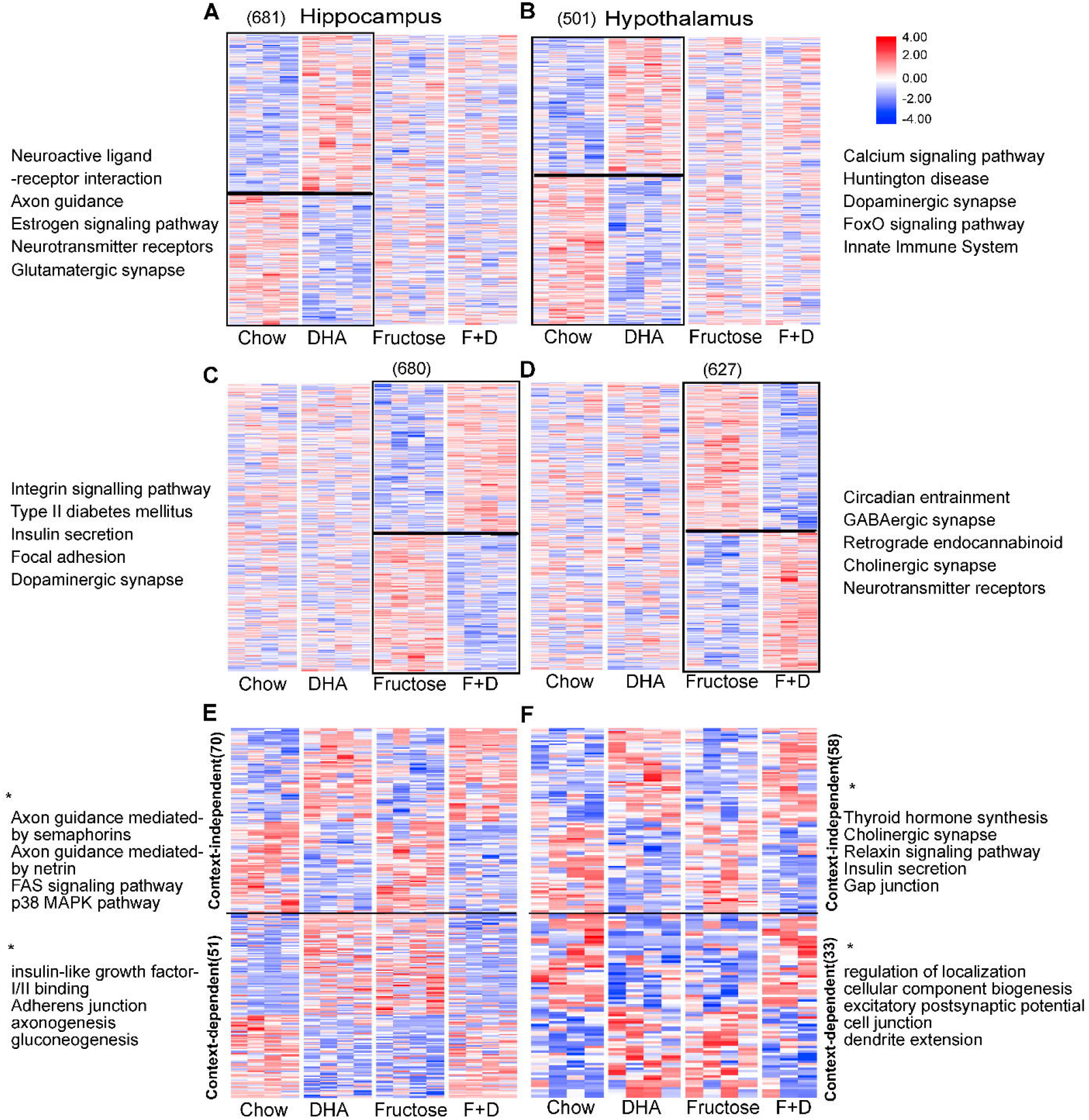
Heatmap of methylation changes and enriched pathways under DHA supplementation with or without fructose treatment in hippocampus and hypothalamus. DHA DMLs (A, B) and fructose DMLs (C, D) were used to compare the methylation changes between different groups in two brain regions. DMLs shared by DHA vs Ctrl DMLs and fructose vs F + D (fructose + DHA) DMLs were separated into context-dependent/independent categories. Blue to red colors indicate low to high expression values. The genes were mapped within 10 kb from DML. * indicted pathways or GO terms without reaching significant enrichment. Number in brackets represented number of genes.

Functional annotation of the genes adjacent to the DMLs revealed a broad range of pathways from ECM, neuronal signaling, and metabolic pathways to immune and endocrine pathways. Some of these pathways overlapped with those revealed through the transcriptome analysis but unique pathways such as thyroid hormone synthesis was also identified (Figure 4A-D).

### 3.6. Relationship between DMLs and DEGs

To investigate the relationship between DNA methylation and gene expression, we mapped the DNA methylation loci to adjacent genes within 10kb distance. Only 14 (7%) and 35 (6%) DHA DEGs identified under the chow diet condition and the fructose condition, respectively, were within 10kb distance to the DMLs in hippocampus; 17 (3%) and 39 (4%) DHA DEGs in the hypothalamus under chow and fructose diet condition, respectively (Table 2). A subset of these genes were shown to be significantly correlated with DMLs (Figure 5). The methylation within *Taok3* is negatively associated with obesity in children ^[22]^. *Nptx2*, a member of neuronal pentraxins, has be linked to hippocampal synaptic plasticity in developmental and adult mice ^[23]^. These results suggest a limited direct correlation between DMLs and DEGs measured at the same time point.

**Table 2.**
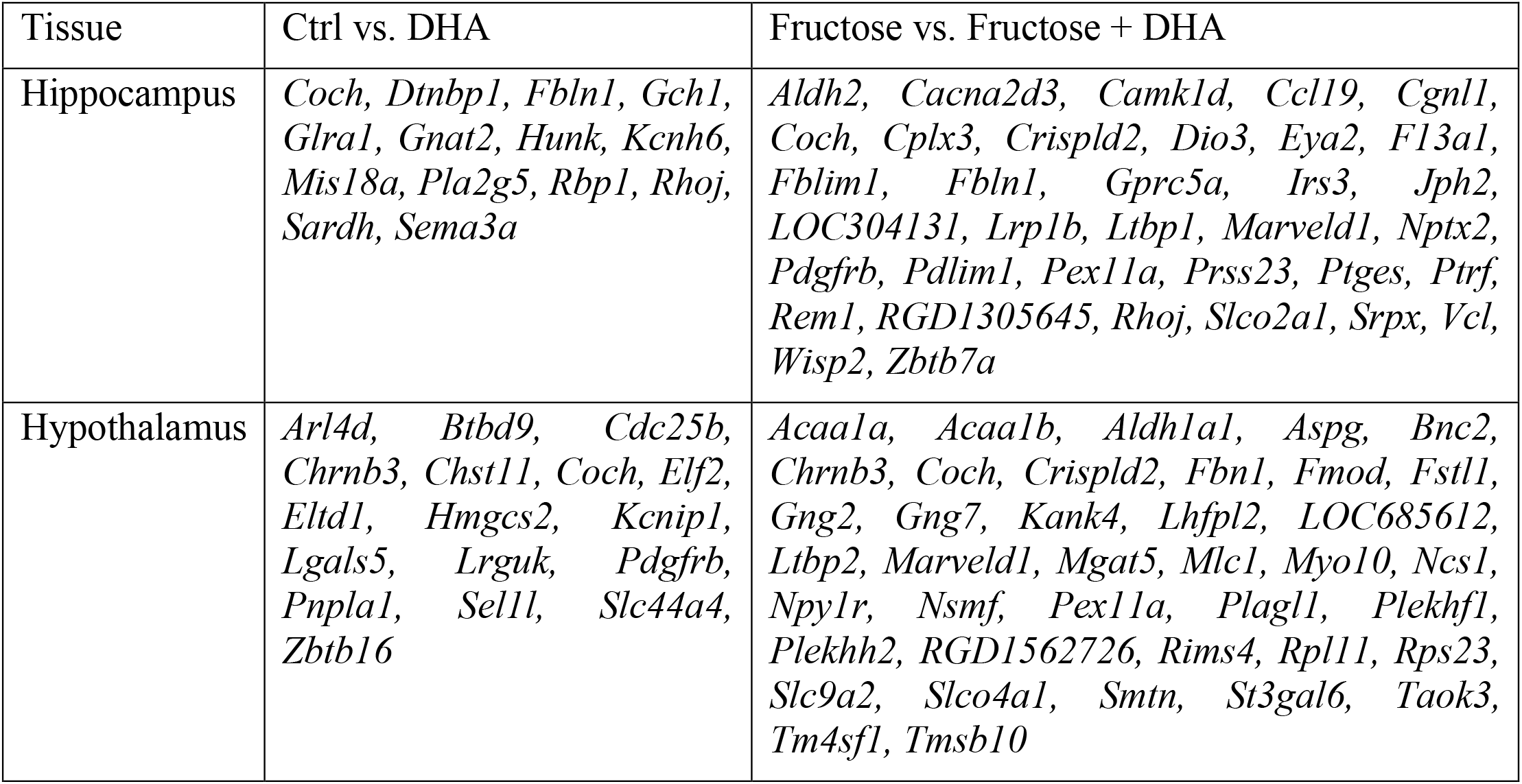
DEGs within a 10kb distance of DMLs affected by DHA on a chow diet vs fructose diet background.

**Figure 5.**
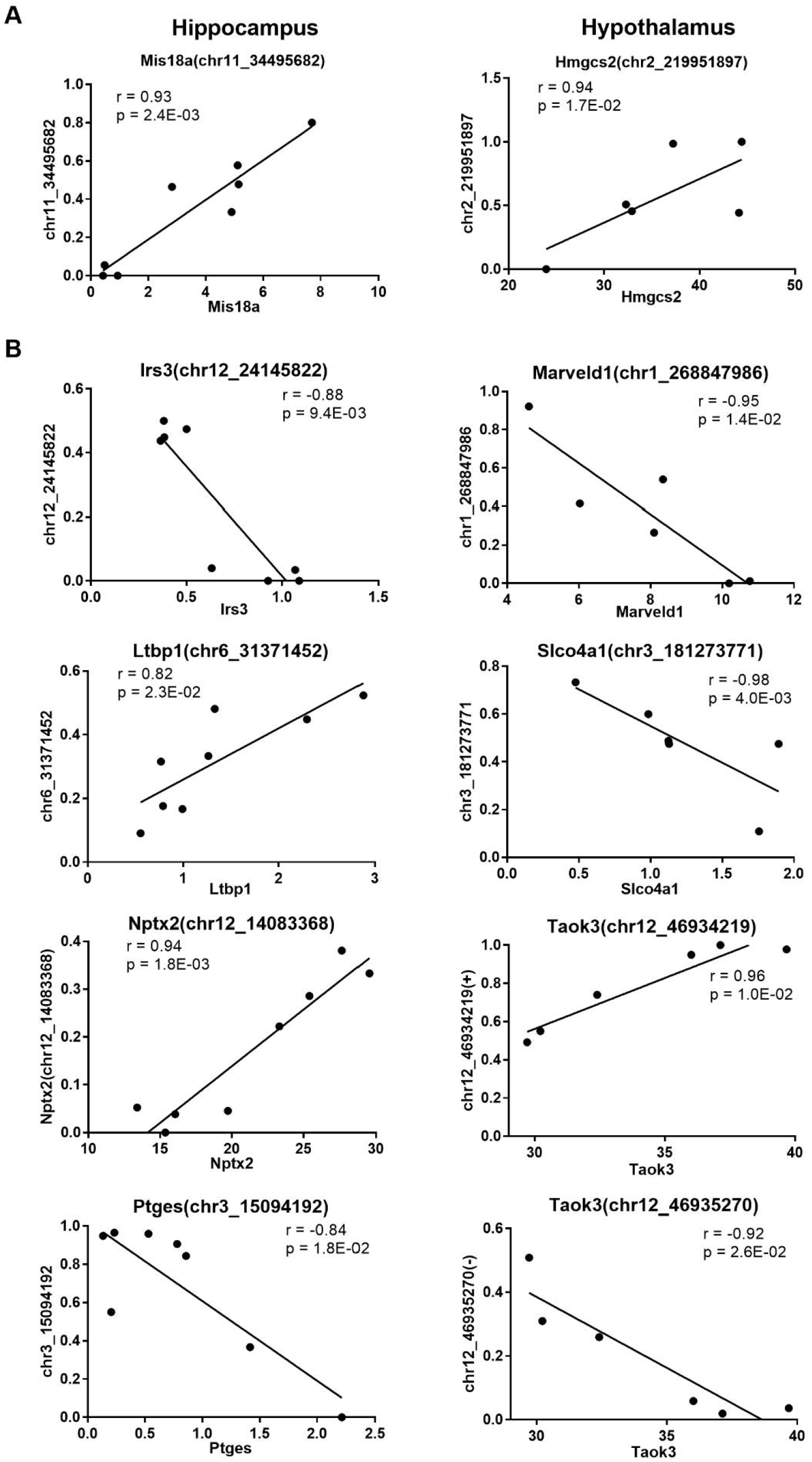
Correlation between DNA methylation and gene expression under chow (A) and fructose (B) background. The p-value was determined using Pearson correlation test and corrected using Benjamini–Hochberg procedure. +: sense strand; -: anti-sense strand.

Some interesting genes among DEGs with local DMLs (Table 2) are transcription factors *Zbtb16* in hypothalamus (chow condition) and *Zbtb7a* in hippocampus (fructose condition). *Crispld2*, is required for the control of membrane trafficking during axon development of hippocampal neurons of rats ^[24]^. *Fmod*, encoding the ECM proteoglycan fibromodulin, showed both epigenetic and transcriptomic changes and was a shared KD identified in the above transcriptome network analysis. Epigenetic modification of these regulatory genes upon DHA supplementation could trigger alterations in downstream target genes and networks.

## 4. Discussion

By comparing the multidimensional effects of DHA, encompassing cardiometabolic and cognitive phenotypes and multiomics alterations in two critical brain regions, between a chow diet background and a fructose consumption context, our study shows that dietary DHA supplementation improved select metabolic traits and brain function, and induced transcriptomic and epigenetic alterations in hypothalamic and hippocampal tissues in both context-independent and context-specific manners.

On the phenotypic level, DHA supplementation significantly reduced serum triglyceride on the fructose diet condition but had a non-significant decreasing trend in the chow condition. This is consistent with previous studies that suggested diets high in long chain omega-3 fatty acids, particularly EPA and DHA, reduces blood triglyceride levels ^[25]^. Similarly, in terms of glycemic phenotypes, insulin resistance index, and memory retention, DHA did not affect these phenotypes significantly when examined on the chow diet background, but significantly improved these phenotypes in fructose-treated animals ^[6]^. These context-specific effects observed in our rat model agree with the findings from a previous human meta-analysis study, which revealed that fish oil supplementation had no effects on insulin sensitivity when all individuals were considered but had beneficial effects on insulin sensitivity among individuals with at least one symptom of metabolic disorders in subgroup analysis ^[26]^. These results indicate that the beneficial effects of DHA on metabolism and cognition need to be considered in the context of the pathophysiological states of individuals.

To explore the mechanisms underlying the context-dependent and independent phenotypic effects of DHA, we examined the transcriptome and epigenome of two brain regions relevant to cognition (hippocampus) and metabolic control (hypothalamus). Pathway analysis of the omics alterations revealed both shared and differential responses in the two brain regions in different metabolic contexts. In particular, genes and pathways related with tissue structure such as Focal adhesion and ECM-receptor interaction, and signal transduction pathways such as PI3K-Akt and Wnt signaling pathways were affected by DHA regardless of the dietary context, although the direction of changes in these genes/pathways are not necessarily the same between contexts. These pathways were also previously observed in PUFA fed rat hypothalamic region ^[27]^, and may represent the core functions of DHA in maintaining cell membrane function and cell signaling.

In the physiological context (chow diet condition), we found DHA modulates hippocampal pathways related with neuronal function such as serotonergic synapse, cholinergic synapse, GABAergic synapse, dopaminergic synapse, glutamatergic synapse, and lipid metabolism. Alterations of similar pathways were also observed in the whole brain of rats fed with fish oil ^[28]^, in the hippocampus of mice on a fish oil diet ^[29]^, and in the hippocampus of stressed mice with n-3 PUFA supplementation ^[30]^. DHA also affected the mTOR signaling pathway in hippocampus. In the hypothalamus, DHA-altered pathways were more related to the innate immunity, such as cytokine-cytokine receptors, NF-kappa B signaling pathway, Toll-like receptor signaling pathway.

In stark contrast, under the condition of fructose-induced metabolic syndrome, DHA altered different sets of pathways in these brain regions: overall downregulation of genes in immune pathways such as cytokine and TGF-beta signaling in hippocampus and upregulation of energy metabolism pathways such as oxidative phosphorylation in hypothalamus (Figure 6). In addition, we found that the thyroid hormone synthesis pathway was uniquely altered by DHA in the hippocampus under fructose consumption. Previously, transthyretin, a thyroid hormone transporter, was found to be enhanced by DHA-rich diet in old rat hippocampus ^[31]^, although fructose was not used in this study.

**Figure 6.**
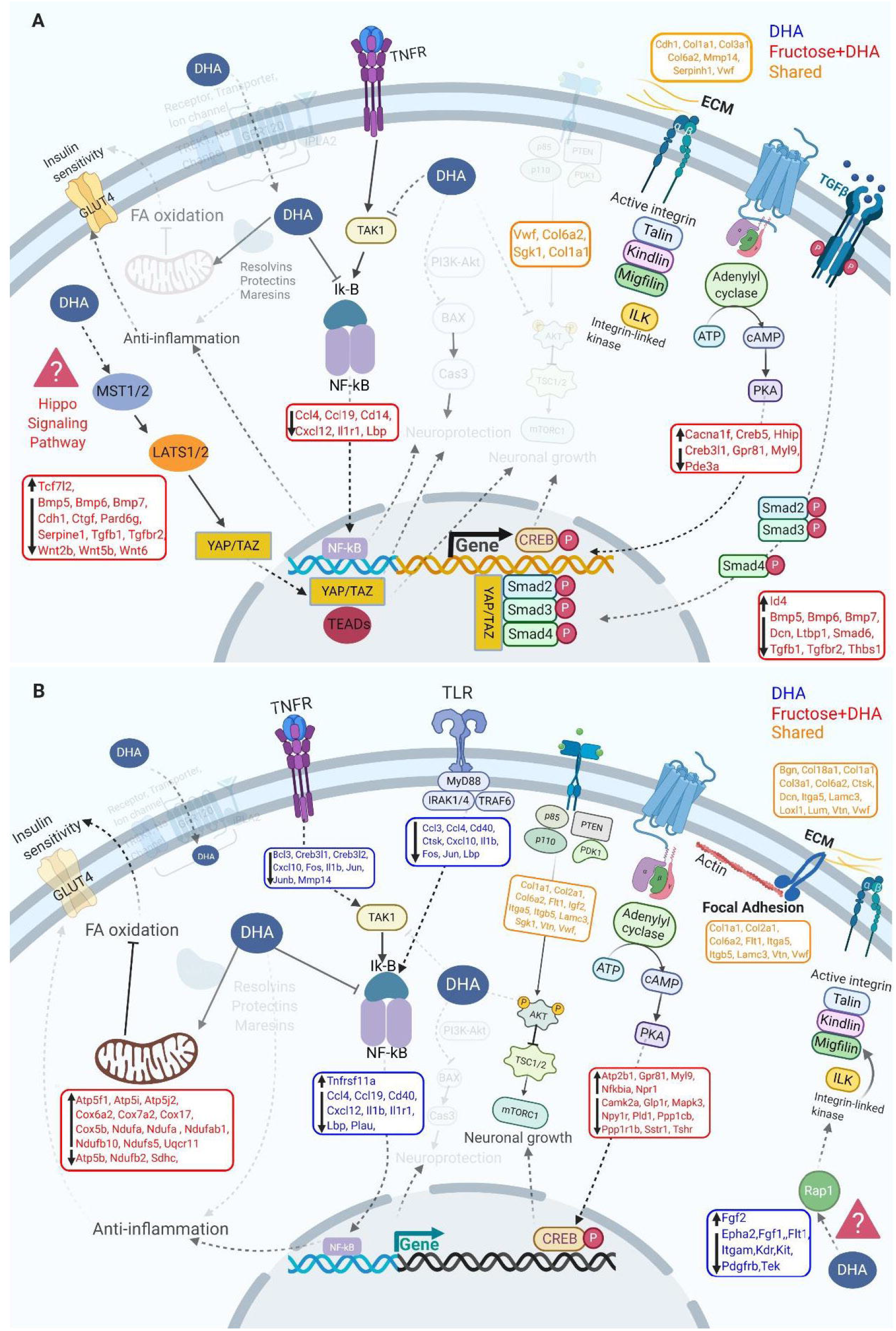
Schematic diagrams of mechanism of action of DHA. The related significantly enriched pathways were highlighted with genes involved in hippocampus (A) and hypothalamus (B).

Between the two dietary conditions (chow vs. fructose) and between tissues, we also identified a consistent set of pathways affected by DHA, including ECM-receptor interaction, Focal adhesion, and PI3K-Akt signaling. These pathways act in or through plasma membrane, which was revealed to be the location of proteins encoded by the majority of genes affected by a fish oil enriched diet ^[32]^. ECM, once known as a pure scaffold to support surrounding cells, is being recognized as an important regulator in neural development, such as proliferation, differentiation, morphogenesis, neuronal migration, formation of axonal process, the myelin sheath, and synapse ^[33]^. Focal adhesion molecules regulate neuronal hyperactivity through the interaction between astrocytes and synapses ^[34]^. Integrin-based focal adhesions connect the ECM to the actin cytoskeleton, which facilitates cell migration and sensing extracellular biochemical and mechanical status ^[35]^ (Figure 6). PI3k-Akt signaling is involved in insulin sensitivity through interaction with the insulin receptor and substrate IRS1/2 in the hypothalamus of rat ^[36]^.

Among the pathways discussed above, some have been previously connected with DHA mode of action, such as mTOR signaling uniquely found in chow condition, and TGF-beta, NF-kappa B, and cAMP signaling pathways specifically enriched under fructose condition in hippocampus (Figure 6A, Table S2). In hypothalamus, the previously implicated pathways included NF-kappa B, Toll-like receptor, and TNF signaling pathways under chow condition, as well as oxidative phosphorylation and cAMP signaling pathway under fructose condition (Figure 6B, Table S2).

In addition to confirming several previously known pathways, our analysis revealed novel pathways affected by DHA, such as the Hippo signaling pathway. Hippo signaling pathway is known to be closely associated with control of organ size and development. Recent studies implicate its role in neural functions, such as neuroinflammation and neuronal cell death ^[37]^. In hypothalamus, Rap1 signaling pathway was also captured, which is involved in neuronal connectivity ^[38]^, hypothalamic inflammation and leptin sensitivity ^[39]^.

Agreeing with the transcriptome-level findings, the DNA methylation loci affected by DHA under the two dietary conditions were also drastically different. These major shifts in the tissue-specific epigenetic loci, genes, and pathways associated with DHA consumption under physiological (chow) and pathological (fructose) conditions highlight the interactions between DHA and the host metabolic states. It is likely that under physiological conditions, DHA balances hippocampal neuronal functions and hypothalamic immune homeostasis. However, in a metabolically challenged state such as high fructose consumption which induces neuroinflammation in hippocampus and perturbs metabolic functions in hypothalamus, DHA mitigates the immune dysfunction induced by fructose in the hippocampus but promotes metabolic homeostasis in the hypothalamus.

Our study also revealed potential network regulators of the genes and pathways affected by DHA within and between contexts. More network regulators were affected by DHA when metabolically challenged with fructose, implicating a much larger network level impact of DHA in this pathological condition. Regardless of conditions, there were many shared network regulators such as *Fmod* and *Bgn*, which were also previously identified as key intermediators of the fructose effects ^[6]^. *Fmod* also showed local DNA methylation changes under DHA treatment (Table 2). Both *Fmod* and *Bgn* encode ECM proteoglycans, which not only provide structural support but also play key regulatory roles in maintaining neuronal functions ^[33, 40]^ and metabolic homeostasis ^[6]^. Among the KDs unique to the chow condition, *Dusp1* is a KD for both the hypothalamic and hippocampal DEGs altered by DHA. *Dusp1* has been implicated in neuroprotection ^[41, 42]^ and has been linked with diabetes related cognitive impairment ^[43]^. *Klf4* is involved in neurite growth and regeneration in hippocampal and cortical neurons, and its dysregulation has been linked to various neurological disorders ^[44, 45]^.

The hippocampal KDs of DHA DEGs unique to the fructose condition include ECM-related genes such as *Lum* encoding Lumican, a protein implicated in collagen binding ^[46, 47]^, and the collagen gene *Col1a1*. Among KDs unique to the fructose condition in the hypothalamus is *Hcar1*, which is involved in the modulation of neuronal activity induced by lactate ^[48]^ in response to physical muscle exertion and resultant communication with the brain enhancing cerebral angiogenesis ^[49]^. Another hypothalamic KD unique to the fructose condition is *Osr1*, a regulator in GABA-mediated depolarization process in brain development ^[50]^.

By employing multidimensional approaches to investigate DHA effects on different tissues under physiological condition and diseased condition, our comprehensive study reveals the context-specific activities of DHA and the underlying molecular mechanism in the form of pathways and regulatory networks. Some of the key regulators uncovered, such as *Fmod* an *Bgn*, have been experimentally validated in terms of their effects on the network genes, cognition and metabolism ^[51, 52]^. However, it is of importance to point out some potential limitations of this study. We acknowledge that only fructose was selected as an unhealthy diet in the current study, yet other types of sugars or combinations with other nutrients need to be tested. Secondly, individual genes or loci from our high throughput omics studies may need further replication, although we note that previous studies have demonstrated the accuracy of high throughput technologies ^[53-56]^.

In summary, DHA exerts distinct influence on metabolic traits and cognition between physiological and pathological conditions. Further investigation revealed transcriptomic and methylome changes in response to DHA under two conditions, offering mechanistic insights into the context-dependent pathways, networks, and key regulators, which may contribute to the differential metabolic and cognitive responses displayed. Our findings offer molecular support of the need for context-specific investigation of PUFAs to facilitate precision nutrition.

## Acknowledgments

X.Y. and F.G.P. are supported by R01 DK104363.

## Author Contributions

X.Y. and F.G.P. designed research; Q.M. conducted omics profiling experiments and A.R. conducted rat experiments; G.Z., Q.M. and M.B. analyzed data; G.Z., M.B., and X.Y. wrote the paper. X.Y. has primary responsibility for final content. All authors read and approved the final manuscript.

## Abbreviations

PUFAs: Long chain polyunsaturated fatty acids
EPA: Eicosapentaenoic acid
T2D: Type 2 diabetes
CHD: Coronary heart disease
CVD: Cardiovascular disease
RNA-Seq: RNA Sequencing
FDR: False discovery rate
GEO: Gene Expression Omnibus
wKDA: weighted key driver analysis
RRBS: Reduced Representation Bisulfite Sequencing
DMLs: Differentially methylated loci

